# Integrated sample inactivation, amplification, and Cas13-based detection of SARS-CoV-2

**DOI:** 10.1101/2020.05.28.119131

**Authors:** Jon Arizti-Sanz, Catherine A. Freije, Alexandra C. Stanton, Chloe K. Boehm, Brittany A. Petros, Sameed Siddiqui, Bennett M. Shaw, Gordon Adams, Tinna-Solveig F. Kosoko-Thoroddsen, Molly E. Kemball, Robin Gross, Loni Wronka, Katie Caviness, Lisa E. Hensley, Nicholas H. Bergman, Bronwyn L. MacInnis, Jacob E. Lemieux, Pardis C. Sabeti, Cameron Myhrvold

## Abstract

The COVID-19 pandemic has highlighted that new diagnostic technologies are essential for controlling disease transmission. Here, we develop SHINE (SHERLOCK and HUDSON Integration to Navigate Epidemics), a sensitive and specific integrated diagnostic tool that can detect SARS-CoV-2 RNA from unextracted samples. We combine the steps of SHERLOCK into a single-step reaction and optimize HUDSON to accelerate viral inactivation in nasopharyngeal swabs and saliva. SHINE’s results can be visualized with an in-tube fluorescent readout — reducing contamination risk as amplification reaction tubes remain sealed — and interpreted by a companion smartphone application. We validate SHINE on 50 nasopharyngeal patient samples, demonstrating 90% sensitivity and 100% specificity compared to RT-PCR with a sample-to-answer time of 50 minutes. SHINE has the potential to be used outside of hospitals and clinical laboratories, greatly enhancing diagnostic capabilities.

## Introduction

Point-of-care diagnostic testing is essential to prevent and effectively respond to infectious disease outbreaks. Insufficient nucleic acid diagnostic testing infrastructure (*1*) and the prevalence of asymptomatic transmission (*2, 3*) have accelerated the global spread of severe acute respiratory syndrome coronavirus 2 (SARS-CoV-2) (*4*–*6*), with confirmed case counts surpassing 5 million (*7*). Ubiquitous nucleic acid testing — whether in doctor’s offices, pharmacies, or mobile/drive-thru/pop-up testing sites — would increase diagnostic access and is essential for safely reopening businesses, schools, and country borders. Easy-to-use, scalable diagnostics with a quick turnaround time and limited equipment requirements would fulfill this major need and have the potential to alter the trajectory of this global pandemic.

The paradigm for nucleic acid diagnostic testing is a centralized model where patient samples are sent to large clinical laboratories for processing and analysis. RT-qPCR, the highly specific and sensitive current gold-standard for SARS-CoV-2 diagnosis (*8*), requires laboratory infrastructure for nucleic acid extraction, thermal cycling, and analysis of assay results. The need for thermocyclers can be eliminated through the use of isothermal (*i.e.*, single temperature) amplification methods, such as loop-mediated isothermal amplification (LAMP) and recombinase polymerase amplification (RPA) (*9*–*14*). However, isothermal amplification methods still require technological advances (Qian, Boswell, Chidley, Lu *et al.* submitted) to increase sensitivity on unextracted RNA samples and to reduce non-specific amplification (*15, 16*), which would enable testing at scale outside of laboratories.

Recently developed CRISPR-based diagnostics have the potential to transform infectious disease diagnosis. Both CRISPR-Cas13- and Cas12-based assays have been developed for SARS-CoV-2 detection using extracted nucleic acids as input (*17*–*22*). One such CRISPR-based diagnostic, SHERLOCK (Specific High-sensitivity Enzymatic Reporter unLOCKing), involves two separate steps, starting with extracted nucleic acid: (1) isothermal RPA and (2) T7 transcription and Cas13-mediated collateral cleavage of a single-stranded RNA reporter (*23*) (Fig. 1A). Cas13-based detection is highly programmable and specific, as it relies on complementary base pairing between the target RNA and the CRISPR RNA (crRNA) sequence (*23, 24*). However, in their current state, these technologies require nucleic acid extraction (often using kits that are in short supply) and multiple sample transfer steps, limiting their widespread use. SHERLOCK can be paired with HUDSON (Heating Unextracted Diagnostic Samples to Obliterate Nucleases), which eliminates the need for nucleic acid extraction by using heat and chemical reduction to both destroy RNA-degrading nucleases and lyse viral particles (*25*). Together, SHERLOCK and HUDSON can be performed with limited laboratory infrastructure, solely requiring a heating element. However, the scalability of these methods is currently limited by the need to prepare multiple reaction mixtures and transfer samples between them.

**Fig. 1.**
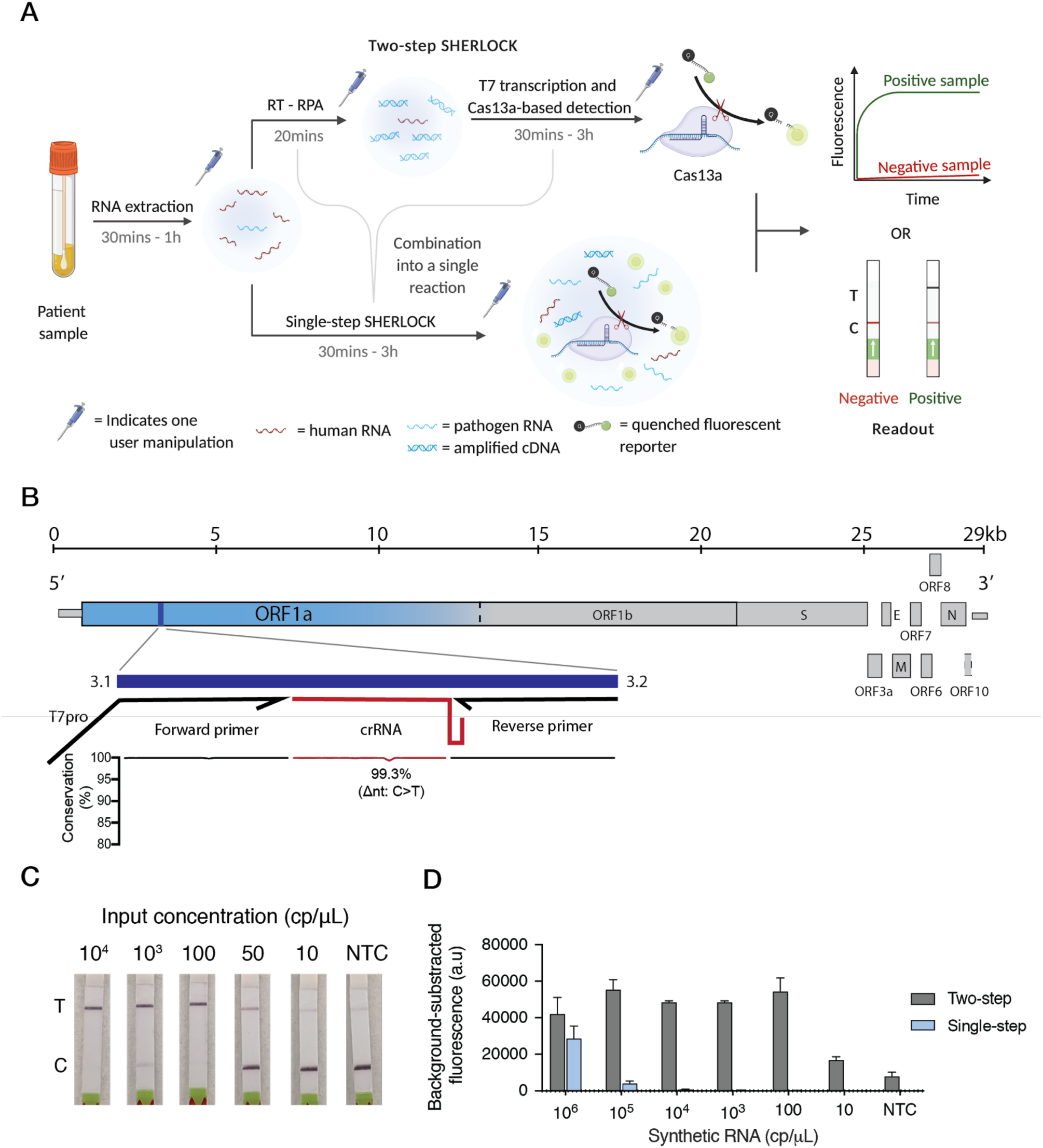
Initial assay development for SHERLOCK-based SARS-CoV-2 detection. (**A**) Schematic of two- and single-step SHERLOCK assays using RNA extracted from patient samples with a fluorescent or colorimetric readout. Times, range of suggested incubation times; pipette, step involving user manipulation; RT-RPA, reverse transcriptase-recombinase polymerase amplification; C, control line; T, test line. (**B**) Schematic of the SARS-CoV-2 genome and SHERLOCK assay location. Sequence conservation across the primer and crRNA binding sites for publicly available SARS-CoV-2 genomes (see Methods for details). Text denotes nucleotide position with lowest percent conservation across the assay location. ORF, open reading frame; narrow rectangles, untranslated regions; dashed border, unlikely to be expressed (*32*). (**C**) Colorimetric detection of synthetic RNA using two-step SHERLOCK after 30 min. NTC_r, non-template control introduced in RPA, NTC_d, non-template control introduced in detection; T, test line; C, control line. (**D**) Background-subtracted fluorescences of the two-step and original single-step SHERLOCK protocols using synthetic SARS-CoV-2 RNA after 3 h. The 1 h timepoint from this experiment is shown in Fig. 2E. NTC, non-template control introduced in RPA. Error bars, s.d. for 2-3 technical replicates.

To address the current limitations of nucleic acid diagnostics, we developed SHINE (SHERLOCK and HUDSON Integration to Navigate Epidemics) for extraction-free, rapid, and sensitive detection of SARS-CoV-2 RNA. We established a SARS-CoV-2 assay (*18*), then combined SHERLOCK’s amplification and Cas13-based detection steps, decreasing user manipulations and assay time (Fig. 1A). We demonstrated that SHINE can detect SARS-CoV-2 RNA in HUDSON-treated patient samples with either a paper-based colorimetric readout, or an in-tube fluorescent readout which can be performed with portable equipment and with reduced risk of sample contamination.

## Results

We first developed a two-step SHERLOCK assay which sensitively detected SARS-CoV-2 RNA at 10 copies per microliter (cp/μL). Using ADAPT, a computational design tool for nucleic acid diagnostics (Metsky *et al. in prep*), we identified a region within open reading frame 1a (ORF1a) of SARS-CoV-2 that comprehensively captures known sequence diversity, with high predicted Cas13 targeting activity and SARS-CoV-2 specificity (Fig. 1B) (*18*). Using both colorimetric and fluorescent readouts, we detected 10 cp/μL of synthetic RNA after incubating samples for 1 h or less, but preparing the reactions required 45-90 minutes of hands-on time depending on the number of samples (Fig. 1C and 1D and fig. S1A). We tested this assay on HUDSON-treated SARS-CoV-2 viral seedstocks, detecting down to 1.31e5 PFU/ml via colorimetric readout (Fig. S1B). Finally, in a side-by-side comparison of our two-step SHERLOCK assay and the CDC RT-qPCR assay, we demonstrated similar limits of detection, reliably identifying 1-10 cp/μL with stochasticity evident at lower viral titers (Fig. S1C).

We sought to develop an integrated, streamlined assay that was significantly less time- and labor-intensive than two-step SHERLOCK. However, when we combined RT-RPA (step 1), T7 transcription, and Cas13-based detection (step 2) into a single step (*i.e.*, single-step SHERLOCK), the sensitivity of the assay decreased dramatically. This decrease was specific for RNA input, and likely due to incompatibility of enzymatic reactions with the given conditions (limit of detection (LOD) 10^6^ cp/μL; Fig. 1D and fig. S2A). As a result, we evaluated whether additional reaction components and optimized reaction conditions could increase the sensitivity and speed of the assay. Addition of RNase H, in the presence of reverse transcriptase, improved the sensitivity of Cas13-based detection of RNA 10-fold (LOD 10^5^ cp/μL; Fig. 2A and fig. S2B and S2C). RNase H likely enhanced the sensitivity by increasing the efficiency of RT through degradation of DNA:RNA hybrid intermediates (Qian, Boswell, Chidley, Lu *et al.* submitted).

**Fig. 2.**
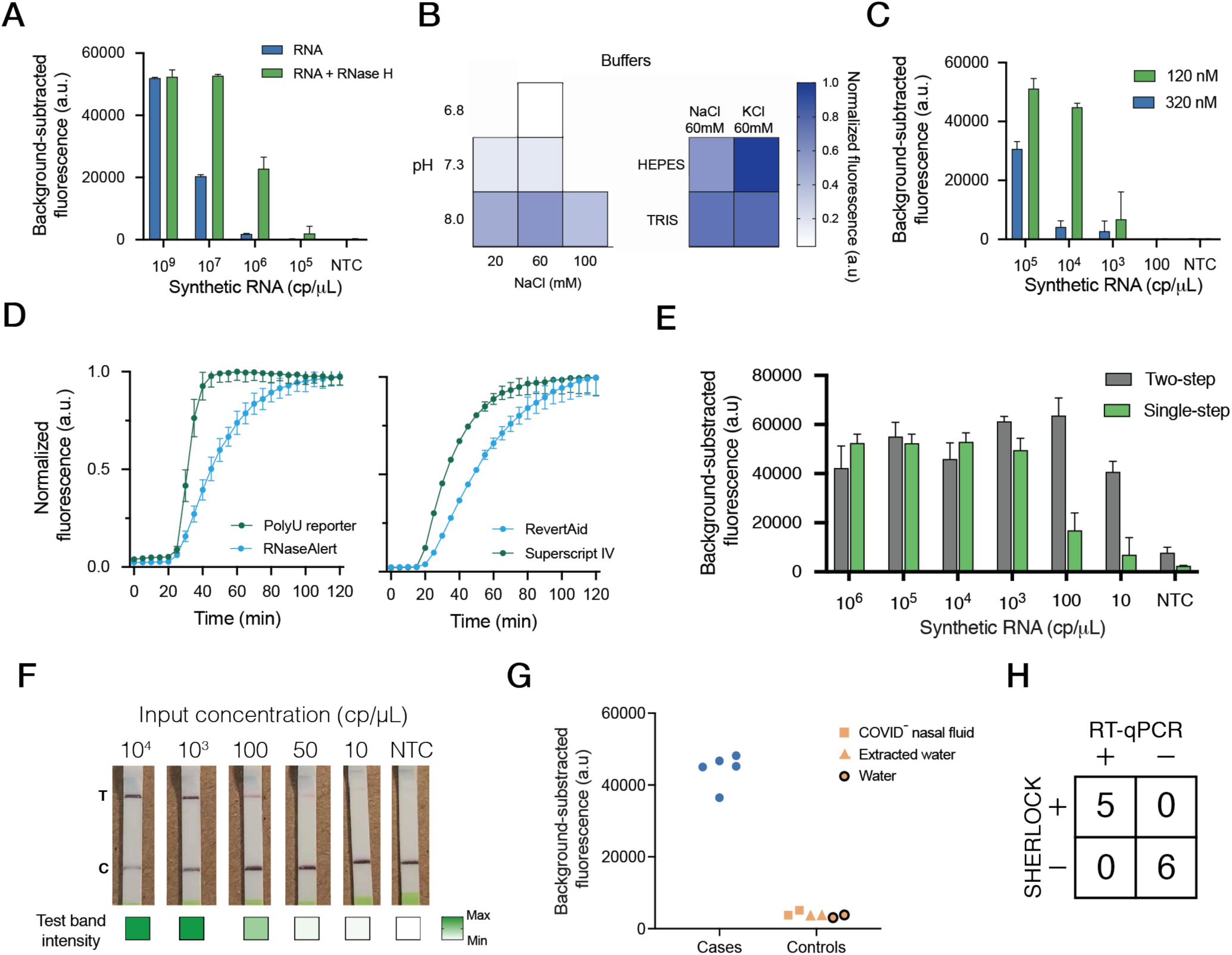
Optimization of the single-step SHERLOCK reaction. (**A**) Background-subtracted fluorescence of Cas13-based detection with synthetic RNA, reverse transcriptase, and RPA primers (but no RPA enzymes) after 3 h. (**B**) Single-step SHERLOCK normalized fluorescence using various buffering conditions after 3 h. (**C**) Background-subtracted fluorescence of single-step SHERLOCK with synthetic RNA and variable RPA forward and reverse primer concentrations after 3 h. (**D**) Single-step SHERLOCK normalized fluorescence over time using two different fluorescent reporters (left) and two different reverse transcriptases (right). (**E**) Background-subtracted fluorescences of the original single-step and optimized single-step SHERLOCK with synthetic RNA after 1 h. Data from the 3 h timepoint from this experiment is shown in Fig. 1D. (**F**) Colorimetric detection of synthetic RNA input using optimized single-step SHERLOCK after 3 h. (**G**) Optimized single-step SHERLOCK background-subtracted fluorescence using RNA extracted from patient samples after 1 h. (**H**) Concordance between SHERLOCK and RT-qPCR for 7 patient samples and 4 controls. For (**C** and **E**), see methods for details about normalized fluorescence calculations. For (**B**,**D**,**F**, and **G**), NTC, non-template control. For (**B**,**D**,**E**, and **F**), error bars, s.d. for 2-3 technical replicates. For (**B** and **D**) RNA input at 10^4^ cp/μL.

Given that each enzyme involved has optimal activity at distinct reaction conditions, we evaluated the role of different pHs, monovalent salt, magnesium, and primer concentrations on assay sensitivity. Optimized buffer, magnesium, and primer conditions resulted in an LOD of 1,000 cp/μL (Fig. 2B and 2C and fig. S2D and S2E). We then improved the speed of Cas13 cleavage and RT to reduce the sample-to-answer time. Given the uracil-cleavage preference of Cas13a (*24, 26, 27*), detection of RNA in the single-step SHERLOCK assay reached half-maximal fluorescence in ∼67% of the time when RNaseAlert was substituted for a polyU reporter (Fig. 2D, left and fig. S3). In addition, reactions containing SuperScript IV reverse transcriptase reached half-maximal fluorescence two times faster than RevertAid (Fig. 2D, right).

Together, these improvements resulted in an optimized single-step SHERLOCK assay that could detect SARS-CoV-2 RNA with reduced sample-to-answer time and equal sensitivity compared to our two-step assay. We quantified the LOD of our optimized single-step SHERLOCK assay on synthetic RNA, detecting as few as 10 cp/μL using a fluorescent readout — 100,000 times more sensitive than the initial assay — and 100 cp/μL using the lateral-flow-based colorimetric readout (Fig. 2E and 2F and fig. S4).

We then evaluated our assay’s performance on SARS-CoV-2 RNA extracted from patient nasopharyngeal (NP) swabs. We compared our fluorescent single-step SHERLOCK assay to previously-performed RT-qPCR using a pilot set of 9 samples. We detected SARS-CoV-2 from 5 of 5 SARS-CoV-2-positive patient samples tested, demonstrating 100% concordance with RT-qPCR, with no false positives for 4 SARS-CoV-2-negative extracted samples nor 2 non-template controls (Fig. 2H and 2I and table S1).

Finally, we paired HUDSON and SHERLOCK with multiple visual readouts to create SHINE (SHERLOCK and HUDSON Integration to Navigate Epidemics), a platform whose results are interpretable by a companion smartphone application (Fig. 3A). In order to reduce total run time, we reduced the incubation time of HUDSON from 30 min to 10 min for both universal viral transport medium (UTM), used for NP swab samples, and for saliva, through the addition of RNase inhibitors (*25*) (Fig. 3B and fig. S5). With this faster HUDSON protocol, we detected 50 cp/μL of synthetic RNA when spiked into UTM and 100 cp/μL when spiked into saliva, using a colorimetric readout (Fig. S6). However, the lateral flow readout requires opening of tubes containing amplified products and interpreting the test band by eye, which introduces risks of sample contamination and user bias, respectively. Thus, we incorporated an in-tube fluorescent readout with SHINE. Within 1 hour, we detected as few as 10 cp/μL of SARS-CoV-2 synthetic RNA in HUDSON-treated UTM and 5 cp/μL in HUDSON-treated saliva with the in-tube fluorescent readout (Fig. 3C and 3D and figs. S7 and S8). To reduce user-bias in interpreting results of this in-tube readout, we developed a companion smartphone app which uses the built-in smartphone camera to image the reaction tubes. The application then calculates the distance of the experimental tube’s pixel intensity distribution from that of a user-selected negative control tube, and returns a binary result indicating the presence or absence of viral RNA in the sample (Fig. 3A and 3E; see Methods for details). Thus, SHINE both minimized equipment requirements and user interpretation bias when implemented with this in-tube readout and the smartphone application.

**Fig. 3.**
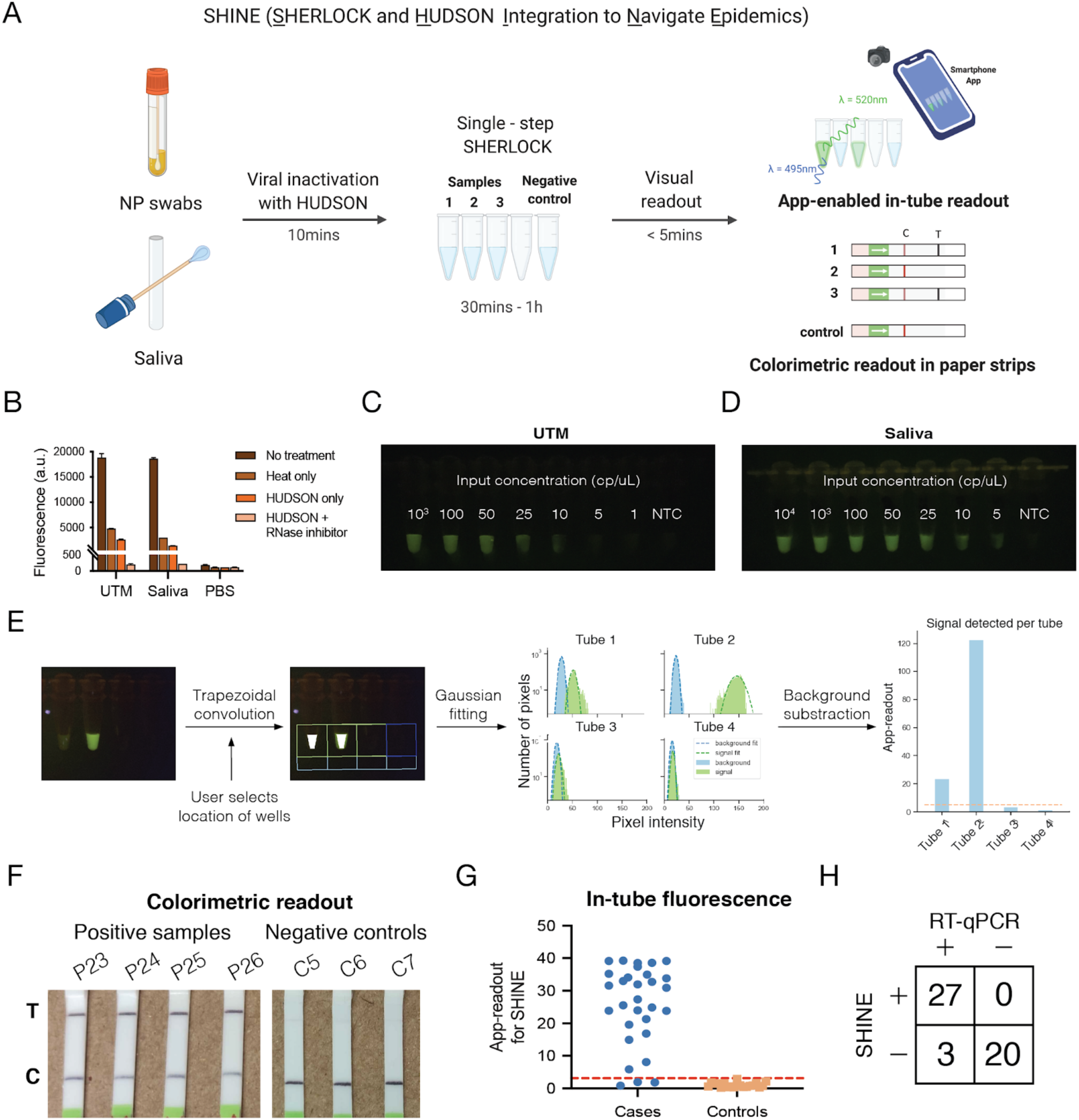
SARS-CoV-2 detection from unextracted samples using SHINE. (**A**) Schematic of SHINE, which is HUDSON paired with single-step SHERLOCK using an in-tube fluorescent or colorimetric readout. Times, range of suggested incubation times; C, control line; T, test line. (**B**) RNaseAlert fluorescence measured after 30 min at room temperature from universal viral transport medium (UTM), saliva, and phosphate buffered saline (PBS) after heat and chemical treatment. (**C**) SARS-CoV-2 RNA detection in HUDSON-treated UTM as measured by single-step SHERLOCK and the in-tube fluorescence readout after 1 h. (**D**) SARS-CoV-2 RNA detection in HUDSON-treated saliva as measured by single-step SHERLOCK and the in-tube fluorescence readout after 1 h. (**E**) Schematic of the companion smartphone application for quantitatively analyzing in-tube fluorescence and reporting binary outcomes of SARS-CoV-2 detection. (**F**) Colorimetric detection of SARS-CoV-2 RNA in unextracted patient NP swabs using the SHINE after 1 h. (**G**) SARS-CoV-2 detection from unextracted patient samples using SHINE and smartphone application quantification of in-tube fluorescence after 40 min. Threshold line determined as average readout value for controls plus 3 standard deviations. (**H**) Concordance table between SHINE and RT-qPCR for 50 patient samples.

We used SHINE to test a set of 50 unextracted, NP samples from 30 RT-qPCR-confirmed, COVID-19-positive patients and 20 COVID-19-negative patients. We used SHINE with the paper-based colorimetric readout on 6 SARS-CoV-2-positive samples and detected SARS-CoV-2 RNA in all 6 positive samples, and in none of the negative controls (100% concordance, Fig. 3F). For all 50 samples, we used SHINE with the in-tube fluorescence readout and companion smartphone application. We detected SARS-CoV-2 RNA in 27 of 30 COVID-19-positive samples (90% sensitivity) and none of the COVID-19-negative samples (100% specificity) after a 10-minute HUDSON and a 40-minute single-step SHERLOCK incubation (Fig. 3G and 3H, fig. S9, and table S1 and S2). Thus, SHINE demonstrated 94% concordance using the in-tube readout with a total run time of 50 minutes. Notably, the RT-qPCR-positive patient NP swabs that SHINE failed to detect tended to have higher Ct values than those that SHINE detected as positive (*p* = 0.0084 via one-sided Wilcoxon rank sum test; Fig. S10). Moreover, this observation could be related to sample degradation and differences in sample processing, as SHINE samples went through additional freeze-thaw cycles and RT-qPCR was performed on extracted and DNase-treated samples.

## Discussion

Here, we have described SHINE, a simple method for detecting viral RNA from unextracted patient samples with minimal equipment requirements. SHINE’s simplicity matches that of the most streamlined nucleic acid diagnostics. Furthermore, the in-tube fluorescence readout and companion smartphone application lends themselves to scalable, high-throughput testing and automated interpretation of results. SHINE’s simplicity and CRISPR-based programmability underscore its potential to address diagnostic needs during the COVID-19 pandemic, and in outbreaks to come.

Additional advances are still required for diagnostic testing to occur in virtually any location. Ideally, all steps would be performed at ambient temperature (without heat), in 15 minutes or less, using a colorimetric readout that does not require tube opening. Existing nucleic acid diagnostics, to our knowledge, are not capable of meeting all these requirements simultaneously. Sample collection without UTM (*i.e.*, “dry swabs”) combined with spin-column-free extraction buffers, and incorporation of solution-based, colorimetric readouts could address these limitations (*28*–*31*). Together, these advances could greatly enhance the accessibility of diagnostic testing and provide an essential tool in the fight against infectious diseases. By reducing personnel time, equipment, and assay time-to-results without sacrificing sensitivity or specificity, we have taken steps towards the development of such a tool.

## Materials and Methods

Detailed information about reagents, including the commercial vendors and stock concentrations, is provided in Table S3.

### Clinical samples and ethics statement

Clinical samples were acquired from clinical studies evaluated and approved by the Institutional Review Board/Ethics Review Committee of the Massachusetts General Hospital and Massachusetts Institute of Technology (MIT). The Office of Research Subject Projection at the Broad Institute of MIT and Harvard University approved use of samples for the work performed in this study.

### Extracted sample preparation and RT-qPCR testing

Nasal swabs were collected and stored in universal viral transport medium (UTM; BD) and stored at -80 °C prior to nucleic acid extraction. Nucleic acid extraction was performed using MagMAX™ *mir*Vana™ Total RNA isolation kit. The starting volume for the extraction was 250 μl and extracted nucleic acid was eluted into 60 μl of nuclease-free water. Extracted nucleic acid was then immediately Turbo DNase-treated (Thermo Fisher Scientific), purified twice with RNACleanXP SPRI beads (Beckman Coulter), and eluted into 15 μl of Ambion Linear Acrylamide (Thermo Fisher Scientific) water (0.8%).

Turbo DNase-treated extracted RNA was then tested for the presence of SARS-CoV-2 RNA using a lab-developed, probe-based RT-qPCR assay based on the N1 target of the CDC assay. RT-qPCR was performed on a 1:3 dilution of the extracted RNA using TaqPath™ 1-Step RT-qPCR Master Mix (Thermo Fisher Scientific) with the following forward and reverse primer sequences, respectively: GACCCCAAAATCAGCGAAAT, TCTGGTTACTGCCAGTTGAATCTG. The RT-PCR assay was run with a double-quenched FAM probe with the following sequence: 5’-FAM-ACCCCGCATTACGTTTGGTGGACC-BHQ1-3’. RT-qPCR was run on a QuantStudio 6 (Applied Biosystems) with RT at 48 °C for 30 min and 45 cycles with a denaturing step at 95 °C for 10 s followed by annealing and elongation steps at 60 °C for 45 s. The data were analyzed using the Standard Curve (SC) module of the Applied Biosystems Analysis Software.

### SARS-CoV-2 assay design and synthetic template information

SARS-CoV-2-specific forward and reverse RPA primers and Cas13-crRNAs were designed as previously described (*18*). In short, the designs were algorithmically selected, targeting 100% of 20 published SARS-CoV-2 genomes, and predicted by a machine learning model to be highly active (Metsky *et al. in prep*). Moreover, the crRNA was selected for its high predicted specificity towards detection of SARS-CoV-2, versus related viruses, including other bat and mammalian coronaviruses and other human respiratory viruses (https://adapt.sabetilab.org/covid-19/).

Synthetic DNA targets with appended upstream T7 promoter sequences (5’-GAAATTAATACGACTCACTATAGGG-3’) were ordered as double-stranded DNA (dsDNA) gene fragments from IDT, and were *in vitro* transcribed to generate synthetic RNA targets. *In vitro* transcription was conducted using the HiScribe T7 High Yield RNA Synthesis Kit (New England Biolabs (NEB)) as previously described (23). In brief, a T7 promoter ssDNA primer (5’-GAAATTAATACGACTCACTATAGGG-3’) was annealed to the dsDNA template and the template was transcribed at 37 °C overnight. Transcribed RNA was then treated with RNase-free DNase I (QIAGEN) to remove any remaining DNA according to the manufacturer’s instructions. Finally, purification occurred using RNAClean SPRI XP beads at 2× transcript volumes in 37.5% isopropanol.

Sequence information for the synthetic targets, RPA primers, and Cas13-crRNA is listed in Table S4.

### Two-step SARS-CoV-2 assay

The two-step SHERLOCK assay was performed as previously described (*18, 23, 25*). Briefly, the assay was performed in two steps: (1) isothermal amplification via recombinase polymerase amplification (RPA) and (2) *Lwa*Cas13a-based detection using a single-stranded RNA (ssRNA) fluorescent reporter. For RPA, the TwistAmp Basic Kit (TwistDx) was used as previously described (*i.e.*, with RPA forward and reverse primer concentrations of 400 nM and a magnesium acetate concentration of 14 mM) (*25*) with the following modifications: RevertAid reverse transcriptase (Thermo Fisher Scientific) and murine RNase inhibitor (NEB) were added at final concentrations of 4 U/µl each, and synthetic RNAs or viral seedstocks were added at known input concentrations making up 10% of the total reaction volume. The RPA reaction was then incubated on the thermocycler for 20 minutes at 41 °C. For the detection step, 1 µl of RPA product was added to 19 µl detection master mix. The detection master mix consisted of the following reagents (final concentrations in master mix listed), with magnesium chloride added last: 45 nM *Lwa*Cas13a protein resuspended in 1× storage buffer (SB: 50 mM Tris pH 7.5, 600 mM NaCl, 5% glycerol, and 2 mM dithiothreitol (DTT); such that the resuspended protein is at 473.7 nM), 22.5 nM crRNA, 125 nM RNaseAlert substrate v2 (Thermo Fisher Scientific), 1× cleavage buffer (CB; 400 mM Tris pH 7.5 and 10 mM DTT), 2 U/µlL murine RNase inhibitor (NEB), 1.5 U/µl NextGen T7 RNA polymerase (Lucigen), 1 mM of each rNTP (NEB), and 9 mM magnesium chloride. Reporter fluorescence kinetics were measured at 37 °C on a Biotek Cytation 5 plate reader using a monochromator (excitation: 485 nm, emission: 520 nm) every 5 minutes for up to 3 hours.

### Single-step SARS-CoV-2 assay optimization

The starting point for optimization of the single-step SHERLOCK assay was generated by combining the essential reaction components of both the RPA and the detection steps in the two-step assay, described above (*23, 25*). Briefly, a master mix was created with final concentrations of 1× original reaction buffer (20 mM HEPES pH 6.8 with 60 mM NaCl, 5% PEG, and 5 µM DTT), 45 nM *Lwa*Cas13a protein resuspended in 1× SB (such that the resuspended protein is at 2.26 µM), 136 nM RNaseAlert substrate v2, 1 U/µl murine RNase inhibitor, 2 mM of each rNTP, 1 U/µl NextGen T7 RNA polymerase, 4 U/µl RevertAid reverse transcriptase, 0.32 µM forward and reverse RPA primers, and 22.5 nM crRNA. The TwistAmp Basic Kit lyophilized reaction components (1 lyophilized pellet per 102 µl final master mix volume) were resuspended using the master mix. After pellet resuspension, cofactors magnesium chloride and magnesium acetate were added at final concentrations of 5 mM and 17 mM, respectively, to complete the reaction.

Master mix and synthetic RNA template were mixed and aliquoted into a 384-well plate in triplicate, with 20 µl per replicate at a ratio of 19:1 master mix:sample. Fluorescence kinetics were measured at 37 °C on a Biotek Cytation 5 or Biotek Synergy H1 plate reader every 5 minutes for 3 hours, as described above. We observed no significant difference in performance between the two plate reader models.

Optimization occurred iteratively, with a single reagent modified in each experiment. The reagent condition (*e.g.*, concentration, vendor, or sequence) that produced the most optimal results — defined as either a lower limit of detection (LOD) or improved reaction kinetics (*i.e.*, reaction saturates faster) — was incorporated into our protocol. Thus, the protocol used for every future reagent optimization consisted of the most optimal reagent conditions for every reagent tested previously.

For all optimization experiments, the modulated reaction component is described in the figures, associated captions, or associated legends. Across all experiments, the following components of the master mix were held constant: 45 nM *Lwa*Cas13a protein resuspended in 1× SB (such that the resuspended protein is at 2.26 µM), 1 U/µl murine RNase inhibitor, 2 mM of each rNTP, 1 U/µl NextGen T7 RNA polymerase, and 22.5 nM crRNA, and TwistDx RPA TwistAmp Basic Kit lyophilized reaction components (1 lyophilized pellet per 102 µl final master mix volume). In all experiments, the master mix components except for the magnesium cofactor(s) were used to resuspend the lyophilized reaction components, and the magnesium cofactor(s) were added last. All other experimental conditions, which differ among the experiments due to real-time optimization, are detailed in Table S5.

### Single-step SARS-CoV-2 optimized reaction

The optimized reaction (see Supplementary Protocol for exemplary implementation) consists of a master mix with final concentrations of 1× optimized reaction buffer (20 mM HEPES pH 8.0 with 60 mM KCl and 5% PEG), 45 nM *Lwa*Cas13a protein resuspended in 1× SB (such that the resuspended protein is at 2.26 µM), 125 nM polyU [*i.e.*, 6 uracils (6U) or 7 uracils (7U) in length, unless otherwise stated] FAM quenched reporter, 1 U/µl murine RNase inhibitor, 2 mM of each rNTP, 1 U/µl NextGen T7 RNA polymerase, 2 U/µl Invitrogen SuperScript IV (SSIV) reverse transcriptase (Thermo Fisher Scientific), 0.1 U/µl RNase H (NEB), 120 nM forward and reverse RPA primers, and 22.5 nM crRNA. Once the master mix is created, it is used to resuspend the TwistAmp Basic Kit lyophilized reaction components (1 lyophilized pellet per 102 µl final master mix volume). Finally, magnesium acetate is the sole magnesium cofactor, and is added at a final concentration of 14 mM to generate the final master mix.

The sample is added to the complete master mix at a ratio of 1:19 and the fluorescence kinetics are measured at 37 °C using a Biotek Cytation 5 or Biotek Synergy H1 plate reader as described above.

### Visual detection via in-tube fluorescence and via lateral flow strip

Minor modifications were made to the single-step SARS-CoV-2 optimized reaction to visualize the readout via in-tube fluorescence or lateral flow strip.

For in-tube fluorescence, we generated the single-step master mix as described above, except the 7U FAM quenched reporter was used at a concentration of 62.5 nM. The sample was added to the complete master mix at a ratio of 1:19. Samples were incubated at 37 °C and images were collected after 30, 45, 60, 90, 120 and 180 minutes of incubation, with image collection terminating once experimental results were clear. A dark reader transilluminator (DR196 model, Clare Chemical Research) was used to illuminate the tubes.

For lateral-flow readout, we generated the single-step master mix as described above, except we used a biotinylated FAM reporter at a final concentration of 1 µM rather than the quenched polyU FAM reporters. The sample was added to the complete master mix at a ratio of 1:19. After 1-2 hours of incubation at 37 °C, the detection reaction was diluted 1:4 in Milenia HybriDetect Assay Buffer, and the Milenia HybriDetect 1 (TwistDx) lateral flow strip was added. Sample images were collected 5 min following incubation of the strip.

### In-tube fluorescence reader mobile phone application

To enable smartphone-based fluorescence analysis, we designed a companion application. Using the application, the user captures an image of a set of strip tubes illuminated by a transilluminator. The user then identifies regions of interest in the captured image by overlaying a set of pre-drawn boxes onto experimental and control tubes. Image and sample information is then transmitted to a server for analysis. Within each of the user-selected squares, the server models the bottom of each tube as a trapezoid and uses a convolutional kernel to determine the location of maximal signal within each tube, using data from the green channel of the RGB image. The server then identifies the background signal proximal to each tube and fits a Gaussian distribution around the background signal and around the in-tube signal. The difference between the mean pixel intensity of the background signal and the mean pixel intensity of the in-tube signal is then calculated as the background-subtracted fluorescence signal for each tube. To identify experimentally significant fluorescent signals, a score is computed for each experimental tube; this score is equal to the distance between the experimental and control background-subtracted fluorescence divided by the standard deviation of pixel intensities in the control signal. Finally, positive or negative samples are determined based on whether the score is above (positive, +) or below (negative, -) 1.5, a threshold identified empirically.

### HUDSON protocols

HUDSON nuclease and viral inactivation were performed on viral seedstock as previously described with minor modifications to the temperatures and incubation times (25). In short, 100 mM TCEP (Thermo Fisher Scientific) and 1 mM EDTA (Thermo Fisher Scientific) were added to non-extracted viral seedstock and incubated for 20 minutes at 50 °C, followed by 10 minutes at 95 °C. The resulting product was then used as input into the two-step SHERLOCK assay.

The improved HUDSON nuclease and viral inactivation protocol was performed as previously described, with minor modifications (*25*). Briefly, 100 mM TCEP, 1 mM EDTA, and 0.8 U/µl murine RNase inhibitor were added to clinical samples in universal viral transport medium or human saliva (Lee Biosolutions). These samples were incubated for 5 minutes at 40 °C, followed by 5 minutes at 70 °C (or 5 minutes at 95 °C, if saliva). The resulting product was used in the single-step detection assay. In cases where synthetic RNA targets were used, rather than clinical samples (*e.g.*, during reaction optimization), targets were added after the initial heating step (40 °C at 5 minutes). This is meant to recapitulate patient samples, as RNA release occurs after the initial heating step when the temperature is increased and viral particles lyse.

For optimization of nuclease inactivation using HUDSON, only the initial heating step was performed. The products were then mixed 1:1 with 400 mM RNaseAlert substrate v2 in nuclease-free water and incubated at room temperature for 30 minutes before imaging on a transilluminator or measuring reporter fluorescence on a Biotek Synergy H1 [at room temperature using a monochromator (excitation: 485 nm, emission: 520 nm) every 5 minutes for up to 30 minutes]. The specific HUDSON protocol parameters modified are indicated in the figure captions.

### SHINE

The SHINE assay consists of the optimized HUDSON protocol (described above) with the resulting product used as the sample input into our optimized, one-step SHERLOCK protocol (described above).

### Data analysis and schematic generation

Conservation of SARS-CoV-2 sequences across our SHERLOCK assay was determined using publicly available genome sequences via GISAID. Analysis was based on an alignment of 5376 SARS-CoV-2 genomic sequences. Percent conservation was measured at each nucleotide within the RPA primer and Cas13-crRNA binding sites and represents the percentage of genomes that have the consensus base at each nucleotide position.

As described above, fluorescence values are reported as background-subtracted, with the fluorescence value collected before reaction progression (*i.e.*, the latest time at which no change in fluorescence is observed, usually time 0, 5, or 10 minutes) subtracted from the final fluorescence value (3 hours, unless otherwise indicated).

Normalized fluorescence values are calculated using data aggregated from multiple experiments with at least one condition in common. The maximal fluorescence value across all experiments is set to 1, with fluorescence values from the same experiment set as ratios of the maximal fluorescence value. Common conditions across experiments are set to the same normalized value, and that value is propagated to determine the normalized values within an experiment.

The Wilcoxon rank sum test was conducted in MATLAB (MathWorks). Schematics shown in Fig. 1A and Fig. 3A were created using *BioRender.com*. All other schematics were generated in Adobe Illustrator (v24.1.2). Data panels were primarily generated via Prism 8 (GraphPad), except Figure 3E which was generated using Python (version 3.7.2), seaborn (version 0.10.1) and matplotlib (version 3.2.1) (*33, 34*).

## Supporting information

Supplementary Protocol

## Acknowledgements

We would like to thank E. Rosenberg for kindly providing patient samples used in this study; the Harvard Medical School Systems Biology Department for providing additional laboratory space to perform the work; those researchers and laboratories who generously made SARS-CoV-2 sequencing data publicly available to aid in our assay design; members of the Sabeti lab — E. Normandin, K. DeRuff, K. Lagerborg, M. Bauer, M. Rudy, K. Siddle, A. Lin and A. Gladden-Young — for assisting with patient sample collection and processing; H. Metsky, for his contributions to the assay design; M. Springer, the Springer lab, and the Sabeti lab, notably H. Metsky, A. Lin, and N. Welch for their thoughtful discussions and reading of the manuscript.

## Items included in Supplementary Materials

Supplementary Text

Figs. S1 to S10

Tables S1 to S4

References (*35*-*38*)

## Other Supplementary Files

Table S5

Supplementary Protocol

