## Supplementary Material for "Integrated sample inactivation, amplification, and Cas13-based detection of SARS-CoV-2"

[1]Broad Institute of Massachusetts Institute of Technology (MIT) and Harvard, Cambridge, MA, USA. [2]Harvard-MIT Program in Health Sciences and Technology, Cambridge, MA, USA. [3]Program in Virology, Harvard Medical School, Boston, MA, USA. [4]Harvard-MIT MD-PhD Program, Boston, MA, USA. [5]Computational and Systems Biology PhD program, MIT, Cambridge, MA, USA. [6]Department of Medicine, Division of Infectious Diseases, Massachusetts General Hospital, Boston, MA, USA. [7]Integrated Research Facility, Division of Clinical Research, National Institute of Allergy and Infectious Diseases, National Institutes of Health, Frederick, MD, USA. [8]National Biodefense Analysis and Countermeasures Center, Fort Detrick, MD, USA. [9]Harvard T.H. Chan School of Public Health, Boston, MA, USA. [10]Department of Organismic and Evolutionary Biology, Harvard University, Cambridge, MA, USA. [11]Howard Hughes Medical Institute, Chevy Chase, MD, USA. [12]Massachusetts Consortium on Pathogen Readiness, Boston, MA, USA.

* These authors contributed equally.

[+] These authors jointly supervised the work.

**This PDF file includes:**

Supplementary Text
Figs. S1 to S10
Tables S1 to S4
References (*35*-*38*)

**Supplementary Text**

Supplemental Discussion

SHINE's simplicity and compatibility with multiple sample types uniquely highlight its utility for widespread use. Previously developed CRISPR-based detection methods, such as SHERLOCK and DETECTR, are highly sensitive and specific, but assay results require nucleic acid extraction and multiple sample-manipulation steps (*18*, *19*, *23*, *25*, *26*, *29*, *35*). Integration of HUDSON and single-step SHERLOCK eliminates these limitations. HUDSON rapidly inactivated nucleases in both UTM (used for NP swabs) and saliva, illustrating its utility for downstream SARS-CoV-2 detection. Furthermore, less invasive sample collection is essential for routine or daily testing. Therefore, SHINE's compatibility with saliva samples is particularly important as saliva reliably contains SARS-CoV-2 RNA and its collection is less invasive (*36*–*38*). Additionally, the SHERLOCK component of SHINE could be re-designed for detection of other respiratory or saliva-secreted viruses, as well as new viral strains or mutations, underscoring the technology's future potential in shining light on the SARS-CoV-2 pandemic and other outbreaks.

We have shown that SHINE can be used to readily detect SARS-CoV-2 RNA from NP samples without extraction, but further advancements are required to improve sensitivity across varying sample conditions. Notably, SHINE demonstrates perfect concordance with RT-qPCR in our samples with Ct values below 22.5, only exhibiting stochasticity among the lower-titer samples. The association of RT-qPCR Ct value with SHINE's performance suggests that some of the observed non-concordance in test results may be due to assay sensitivity, possibly combined with degradation of sample material in the time or storage conditions between the two assays as the assays were not performed side-by-side. Improvements in isothermal amplification or HUDSON could also make these methods more comparable in performance.

Further optimization of visual readouts is required to deploy SHINE widely. Our current colorimetric readout using paper-based, lateral flow strips has reduced speed compared to the in-tube fluorescent readout. Alternative visual readouts that do not require inserting a paper strip into each sample or measuring specific wavelengths of light would further decrease user and equipment needs (*30*, *31*). These closed-tube visual readouts significantly decrease the risk of contamination across samples of amplified products. Ultimately, we hope to lyophilize SHINE, simplifying distribution and allowing tests to be shelf-stable.

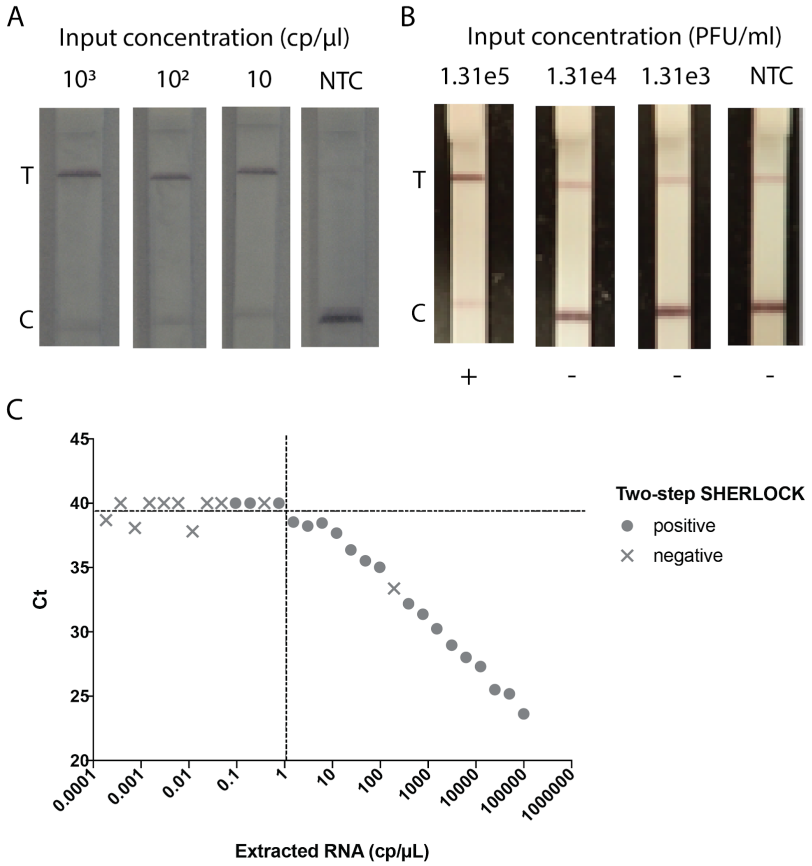


**Fig. S1. Additional two-step SHERLOCK testing.**

(**A**) Colorimetric detection of synthetic DNA using two-step SHERLOCK after 3 h. NTC, non-template control; T, test line; C, control line. (**B**) Colorimetric detection of HUDSON-treated SARS-CoV-2 viral seedstock using two-step SHERLOCK after 3 h. NTC, non-template control; T, test line; C, control line. (**C**) Ct values of real-time RT-qPCR for extracted RNA from SARS-CoV-2 seedstock at various concentrations labelled by the result of our two-step SHERLOCK assay performed side-by-side. The vertical line demarcates 1 cp/µl. The horizontal line demarcates samples with non-quantifiable Ct values (*i.e.* no amplification), imputed as a Ct of 40.

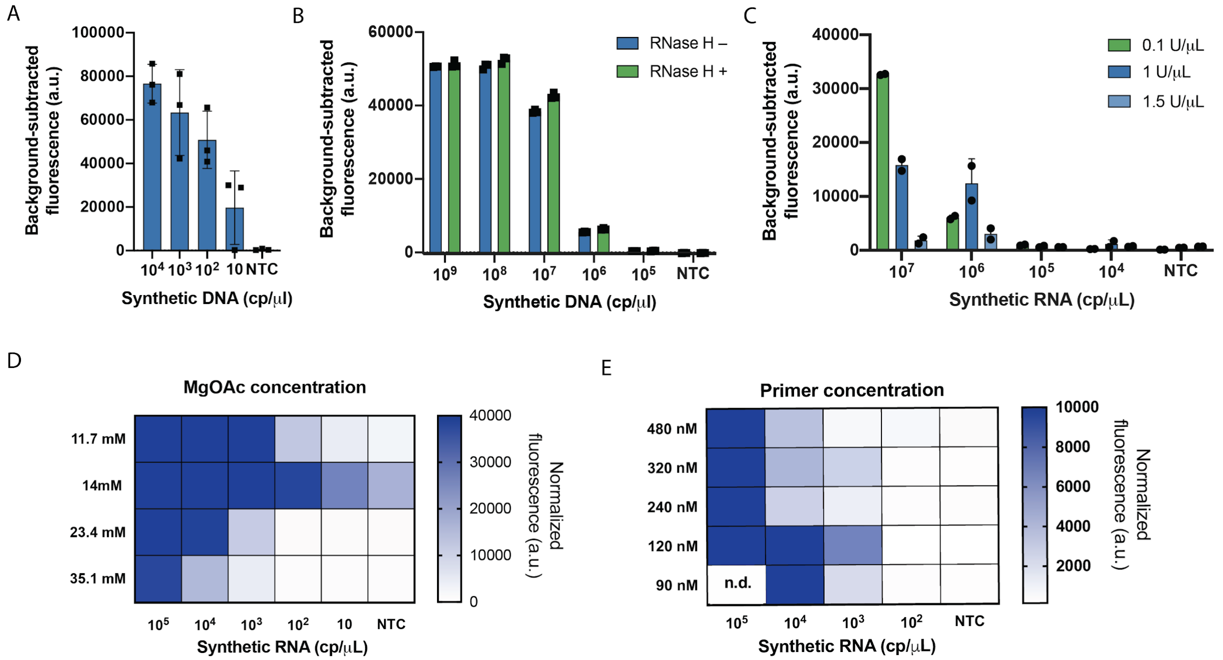


**Fig. S2. Optimization of single-step for improved sensitivity.**

(**A**) Background-subtracted fluorescence detected after the single-step SHERLOCK reaction was incubated for 3 h with DNA as input. (**B**) Background-subtracted fluorescence of the Cas13-detection reaction (no RPA enzymes) with 3 h incubation. RNase H+, final concentration of 0.1 U/µl. (**C**) Background-subtracted fluorescence of the Cas13-detection reaction (no RPA) with 3 h incubation. (**D**) Background-subtracted fluorescence detected after the single-step reaction was incubated for 3 h with varying magnesium concentrations. (**E**) Background-subtracted fluorescence detected after the single-step reaction was incubated for 3 h with varying RPA primer concentrations. For (**A** - **E**), NTC, non-template controls; error bars, s.d. for 2-3 technical replicates. All listed concentrations refer to concentration within the reaction mixture before addition of the oligonucleotide template.

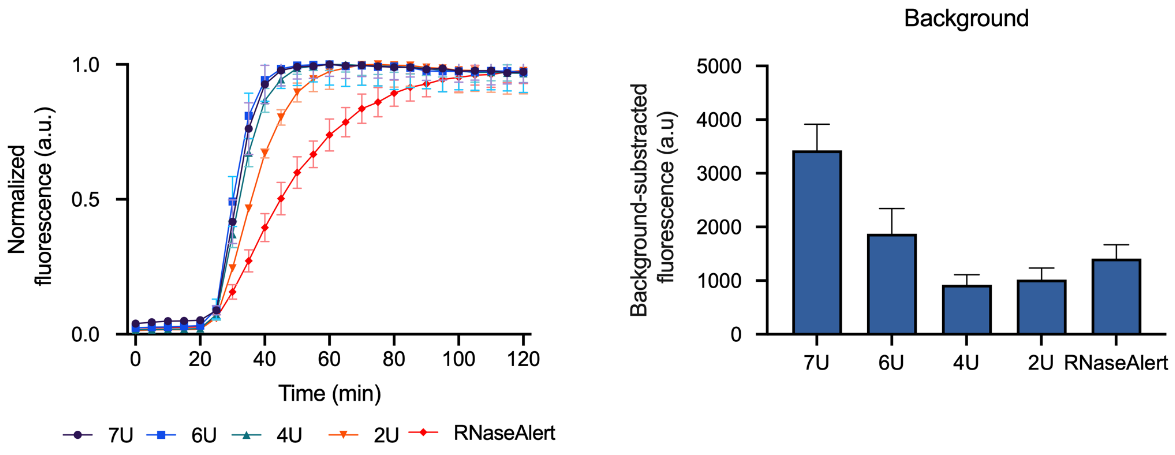


**Fig. S3. Optimization of fluorescent reporter.**

(**A**) Single-step SHERLOCK normalized fluorescence (see methods for details) over time using quenched poly-uracil FAM reporters of varying lengths or RNaseAlert with RNA input at $10^4$ cp/µl. (**B**) Baseline fluorescence of poly-uracil FAM reporters or RNaseAlert in single-step SHERLOCK after 3 h.

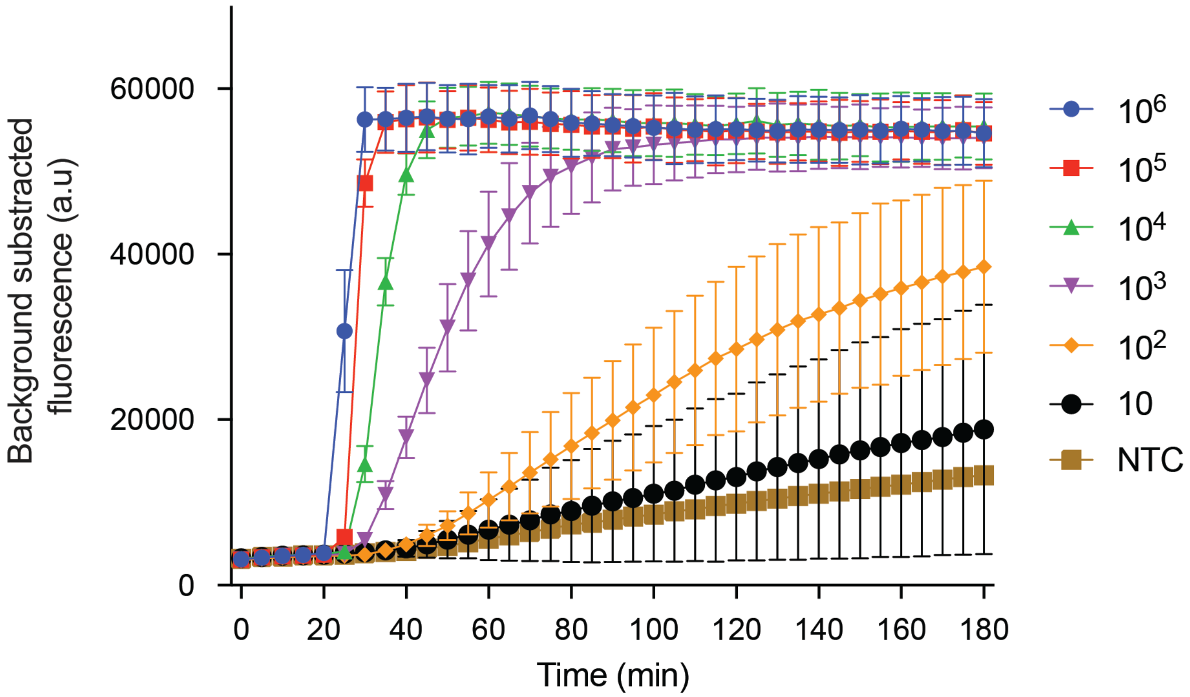


**Fig. S4. Single-step SHERLOCK time course.**

Optimized single-step SHERLOCK assay fluorescence over time at varying RNA input concentrations. Background-subtracted fluorescence at 1h is shown in Fig. 2E. NTC, non-template control; error bars, s.d. for 3 technical replicates. Note: error bars for NTC are present but very small.

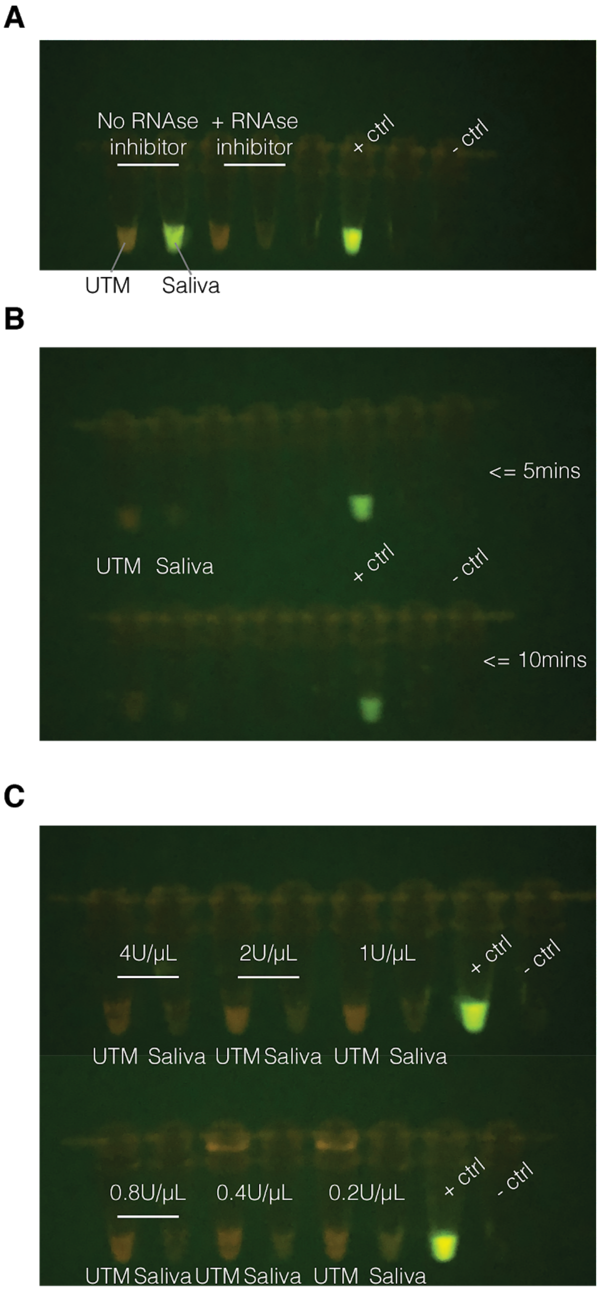


**Fig. S5. HUDSON optimization experiments.**
(**A**) Samples were treated with 100 mM TCEP and 1 mM EDTA and subjected to a 20 min heating step at 50 ˚C. RNase inhibitor, 4 U/µl. (**B**) Samples were treated with 100 mM TCEP, 1mM EDTA, and 4 U/µl RNase inhibitor. (**C**) Samples were treated with 100 mM TCEP and 1 mM EDTA and subjected to a 5 min heating step at 50 ˚C. (**A-C**) Positive and negative controls undergo no treatment. RNaseAlert (final concentration: 200nM) was added immediately after the heating step.

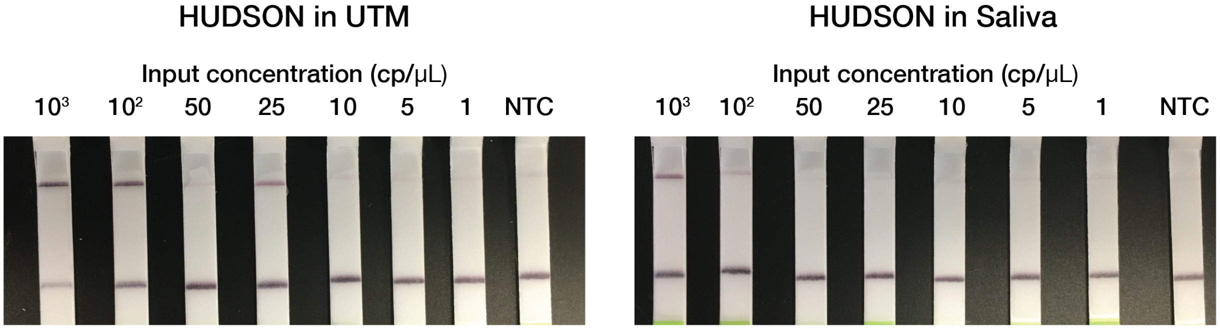


**Fig. S6. SHINE for UTM and saliva with colorimetric detection.**
HUDSON-treated UTM (left) and saliva (right) with synthetic RNA template added after initial heating step. Treated samples were used as input into single-step SHERLOCK with the colorimetric readout.

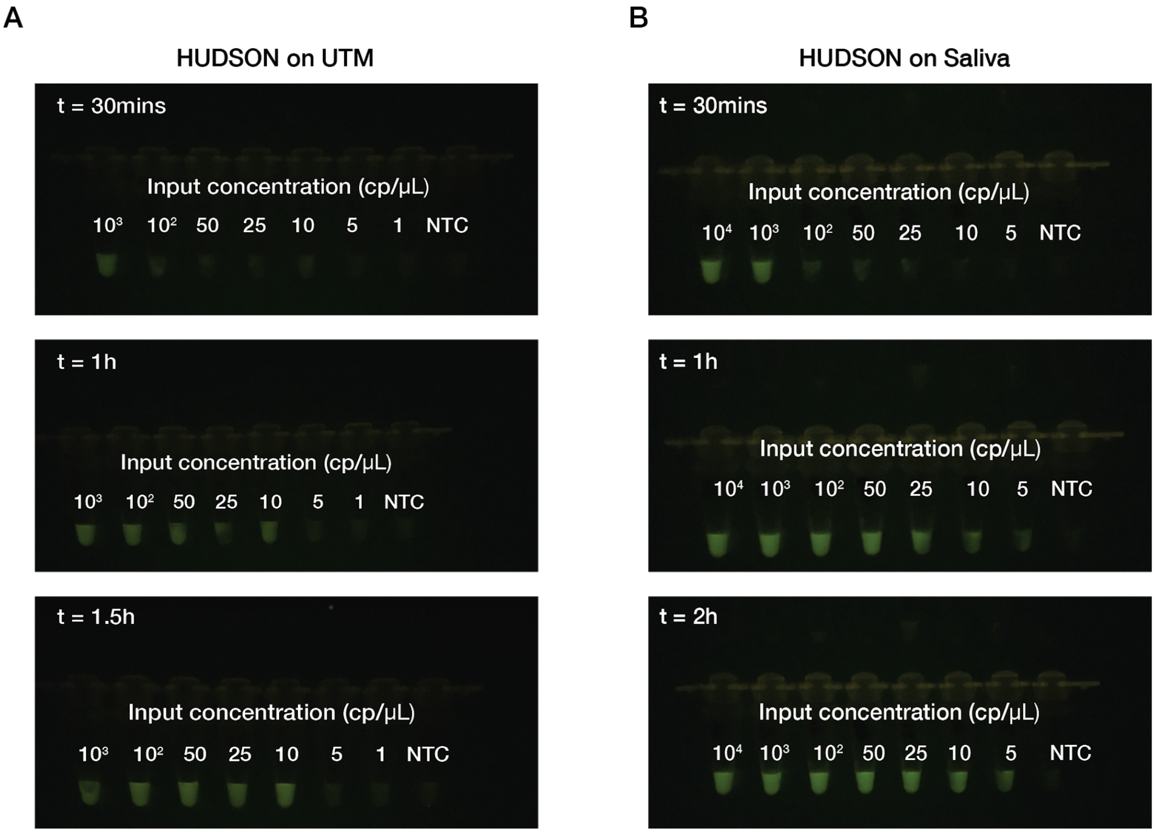


**Fig. S7. SHINE for UTM and saliva with in-tube fluorescent detection.**
(**A** - **B**) HUDSON-treated UTM (**A**) and saliva (**B**) with synthetic RNA template added after initial heating step. Samples were used as input in the single-step SHERLOCK assay. Transilluminator images were captured using a smartphone camera. NTC, non-template control.

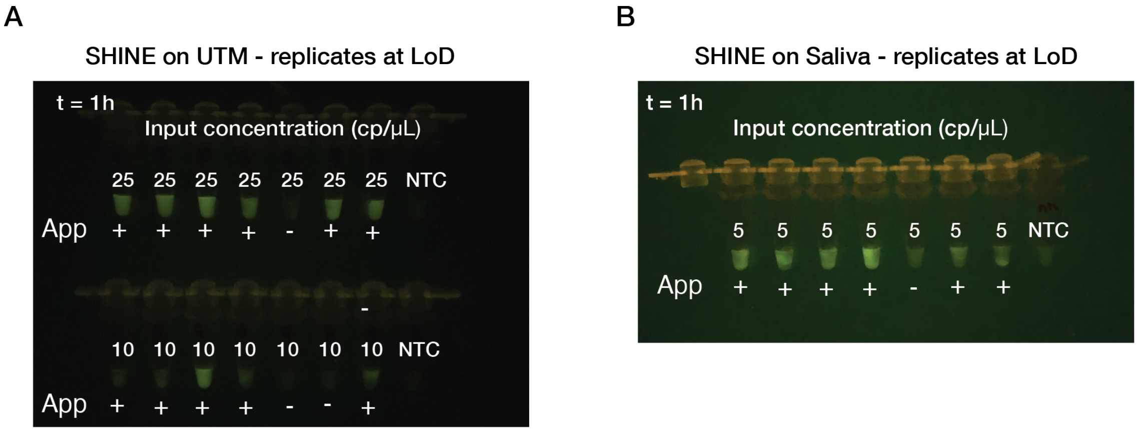


**Fig. S8. Limit of detection of SHINE on UTM and saliva.**
(**A - B**) HUDSON-treated UTM (**A**) and saliva (**B**) with synthetic RNA added after the initial heating step when nucleases are inactivated. Samples were used as input in the single-step SHERLOCK assay incubated for 1 h. Transilluminator images were captured using a smartphone camera and analyzed by the companion smartphone application (App).

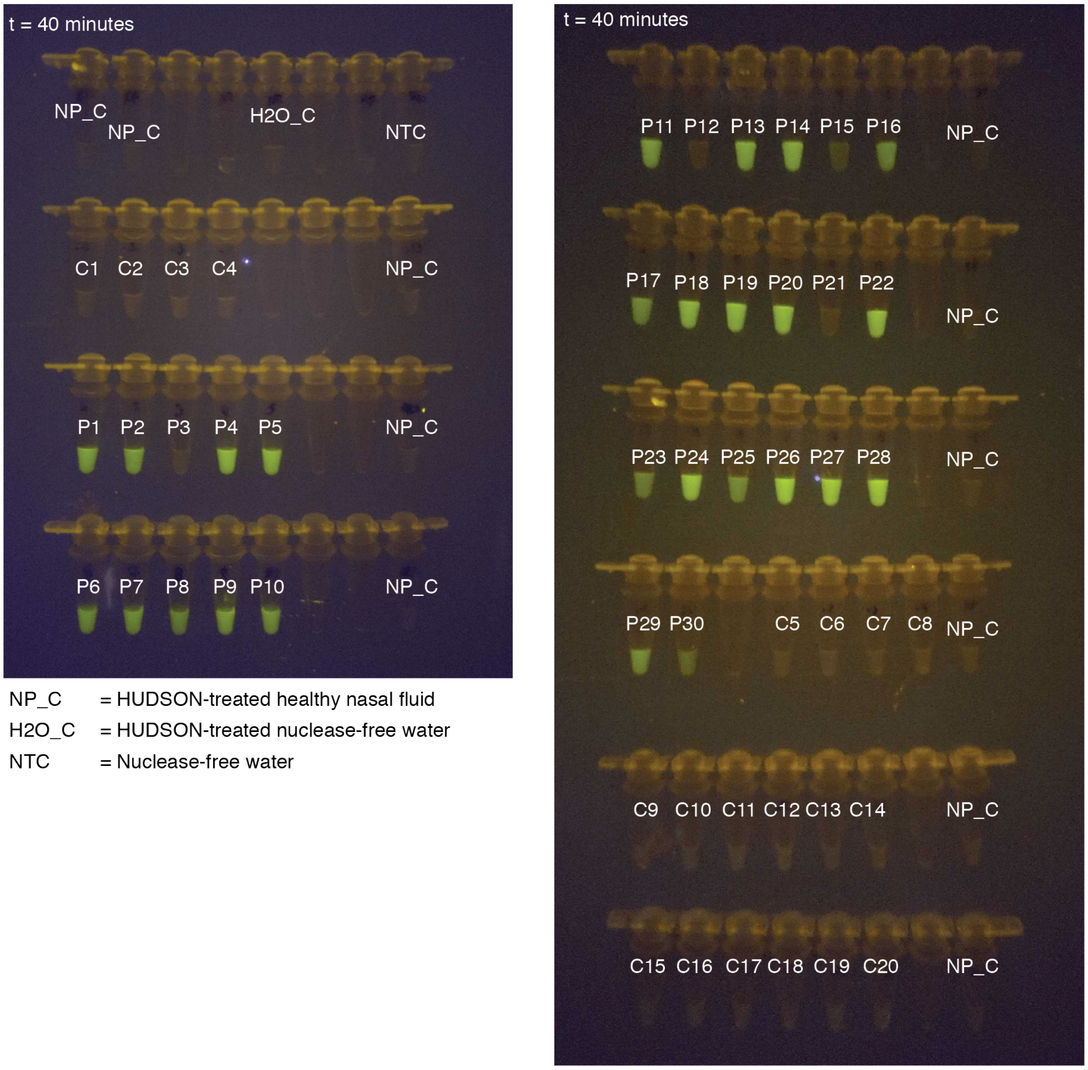


**Fig. S9. SHINE on unextracted patient samples.**
HUDSON-treated NP swabs in UTM were used as input in the single-step SHERLOCK assay. Transilluminator images were captured using a smartphone camera after 40 minutes.

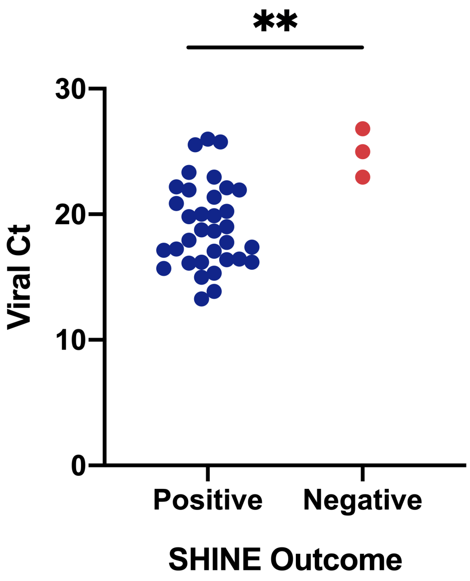


**Fig. S10. SHINE's ability to detect viral RNA is significantly associated with the RT-qPCR threshold cycle.**

Viral Ct values measured by SARS-CoV-2 RT-qPCR of extracted RNA from 30 patient NP samples grouped by the result of SARS-CoV-2 SHINE. The association between viral Ct and SHINE outcome was assessed using a one-sided Wilcoxon rank sum test. **, $p = 0.0084$.

**Table S1: Patient sample information.**

Viral quantity is adjusted as sample was diluted 1:3 for input into in-house RT-qPCR assay. NP, nasopharyngeal swab; UTM, universal viral transport medium.

| Sample name | Sample ID | Sample type | Viral Ct | Adjusted viral quantity (cp/µl) | Figures |
|---|---|---|---|---|---|
| E1 | MA_MGH_00155 | NP in UTM | 25.989 | 892.22 | 2G and H |
| E2 | MA_MGH_00156 | NP in UTM | 17.753 | 85,470.64 | 2G and H |
| E3 | MA_MGH_00157 | NP in UTM | 17.235 | 113,800.532 | 2G and H |
| E4 | MA_MGH_00159 | NP in UTM | 20.861 | 15,298.96 | 2G and H |
| E5 | MA_MGH_00160 | NP in UTM | 20.006 | 24,515.988 | 2G and H |
| E6 | MA_MGH_00166 | NP in UTM | 25.548 | 1,136.696 | 2G and H |
| P1 | MA_MGH_00441 | NP in UTM | 22.109 | 52787.064 | 3G and H |
| P2 | MA_MGH_00442 | NP in UTM | 25.764 | 4386.186 | 3G and H |
| P3 | MA_MGH_00443 | NP in UTM | 26.809 | 2154.246 | 3G and H |
| P4 | MA_MGH_00445 | NP in UTM | 16.429 | 3390658.2 | 3G and H |
| P5 | MA_MGH_00446 | NP in UTM | 17.136 | 2022960.6 | 3G and H |
| P6 | MA_MGH_00447 | NP in UTM | 16.39 | 3426433.56 | 3G and H |
| P7 | MA_MGH_00453 | NP in UTM | 21.935 | 68312.187 | 3G and H |
| P8 | MA_MGH_00454 | NP in UTM | 21.936 | 68293.593 | 3G and H |
| P9 | MA_MGH_00456 | NP in UTM | 15.678 | 5660253.54 | 3G and H |
| P10 | MA_MGH_00458 | NP in UTM | 23.331 | 25530.948 | 3G and H |
| P11 | MA_MGH_00459 | NP in UTM | 15.313 | 7329902.04 | 3G and H |
| P12 | MA_MGH_00460 | NP in UTM | 24.979 | 7977.141 | 3G and H |
| P13 | MA_MGH_00461 | NP in UTM | 16.2 | 3914305.92 | 3G and H |
| P14 | MA_MGH_00463 | NP in UTM | 17.05 | 2157841.26 | 3G and H |
| P15 | MA_MGH_00464 | NP in UTM | 20.234 | 227657.529 | 3G and H |
| P16 | MA_MGH_00465 | NP in UTM | 16.102 | 4201061.4 | 3G and H |
| P17 | MA_MGH_00466 | NP in UTM | 17.933 | 1151605.125 | 3G and H |

| | | | | | |
|---|---|---|---|---|---|
| P18 | MA_MGH_00467 | NP in UTM | 19.023 | 541607.094 | 3G and H |
| P19 | MA_MGH_00468 | NP in UTM | 18.661 | 689269.86 | 3G and H |
| P20 | MA_MGH_00469 | NP in UTM | 14.983 | 9240984 | 3G and H |
| P21 | MA_MGH_00471 | NP in UTM | 22.963 | 33130.494 | 3G and H |
| P22 | MA_MGH_00472 | NP in UTM | 13.271 | 30975183 | 3G and H |
| P23 | MA_MGH_00473 | NP in UTM | 18.757 | 645122.16 | 3F, G and H |
| P24 | MA_MGH_00474 | NP in UTM | 17.397 | 1682047.8 | 3F, G and H |
| P25 | MA_MGH_00475 | NP in UTM | 19.896 | 289419.75 | 3F, G and H |
| P26 | MA_MGH_00476 | NP in UTM | 16.181 | 3970140.3 | 3F, G and H |
| P27 | MA_MGH_00477 | NP in UTM | 19.812 | 305659.512 | 3F, G and H |
| P28 | MA_MGH_00479 | NP in UTM | 13.868 | 20324826 | 3F, G and H |
| P29 | MA_MGH_00480 | NP in UTM | 21.362 | 102417.507 | 3G and H |
| P30 | MA_MGH_00481 | NP in UTM | 22.187 | 57196.188 | 3G and H |

**Table S2: COVID-19 negative patient sample information.**

Control samples were collected prior to the start of the coronavirus pandemic, and therefore are assumed to not contain SARS-CoV-2 RNA.

| Sample name | Sample ID | Sample type | Viral Ct | Adjusted viral quantity (cp/µl) | Figures |
|---|---|---|---|---|---|
| C1 | MA_MGH_00411 | NP in UTM | - | - | 3G and H |
| C2 | MA_MGH_00412 | NP in UTM | - | - | 3G and H |
| C3 | MA_MGH_00413 | NP in UTM | - | - | 3G and H |
| C4 | MA_MGH_00414 | NP in UTM | - | - | 3G and H |
| C5 | MA_MGH_00320 | NP in UTM | - | - | 3F, G and H |
| C6 | MA_MGH_00321 | NP in UTM | - | - | 3F, G and H |
| C7 | MA_MGH_00322 | NP in UTM | - | - | 3F, G and H |
| C8 | MA_MGH_00323 | NP in UTM | - | - | 3F, G and H |
| C9 | MA_MGH_00324 | NP in UTM | - | - | 3G and H |
| C10 | MA_MGH_00325 | NP in UTM | - | - | 3G and H |
| C11 | MA_MGH_00326 | NP in UTM | - | - | 3G and H |
| C12 | MA_MGH_00327 | NP in UTM | - | - | 3G and H |
| C13 | MA_MGH_00328 | NP in UTM | - | - | 3G and H |
| C14 | MA_MGH_00329 | NP in UTM | - | - | 3G and H |
| C15 | MA_MGH_00338 | NP in UTM | - | - | 3G and H |
| C16 | MA_MGH_00353 | NP in UTM | - | - | 3G and H |
| C17 | MA_MGH_00354 | NP in UTM | - | - | 3G and H |
| C18 | MA_MGH_00355 | NP in UTM | - | - | 3G and H |
| C19 | MA_MGH_00356 | NP in UTM | - | - | 3G and H |
| C20 | MA_MGH_00357 | NP in UTM | - | - | 3G and H |

**Table S3: Reagents used in either the optimized one-step assay or in the optimization process.**

HUDSON = Heating Unextracted Diagnostic Samples to Obliterate Nucleases. IVT = *In Vitro* Transcription. PCR = Polymerase Chain Reaction. RPA = Recombinase Polymerase Amplification. SHERLOCK = Specific High Sensitivity Enzymatic Reporter Unlocking.

| Reagent | Reaction | Source | Stock Concentration | Notes |
|---|---|---|---|---|
| EDTA | HUDSON | Thermo Fisher Scientific™ | 0.5 M | |
| TCEP-HCl | HUDSON | Thermo Fisher Scientific™ | 0.5 M | |
| Universal Viral Transport Medium (UTM) | HUDSON | BD | N/A | |
| Saliva, Pooled Human Donors | HUDSON | Lee Biosolutions, Inc. | N/A | |
| RNase Inhibitor | HUDSON; RPA; SHERLOCK | NEB® | 40 U/µL | Murine |
| HiScribe™ T7 High Yield RNA Synthesis Kit | IVT | NEB® | N/A | |
| RNAClean XP | IVT | Beckman Coulter, Inc. | N/A | |
| RNase-Free DNase I | IVT | QIAGEN | | |
| T7 Promoter ssDNA Primer | IVT | Integrated DNA Technologies™ | | *Table S4* for sequences |
| Ambion® Linear Acrylamide | PCR | Thermo Fisher Scientific™ | 5 mg/mL | |
| AMPure XP | PCR | Beckman Coulter, Inc. | N/A | |
| Inter-amplicon Primer | PCR | Integrated DNA Technologies™ | 5 µM | *Table S4* for sequences |
| MagMAX™ *mir*Vana™ Total RNA Isolation Kit | PCR | Thermo Fisher Scientific™ | N/A | |
| PCR Primers (forward, reverse) | PCR | Integrated DNA Technologies™ | | *Table S4* for sequences |

| | | | | |
|---|---|---|---|---|
| Power SYBR® Green Master Mix | PCR | Thermo Fisher Scientific™ | N/A | |
| TaqPath™ 1-Step RT-qPCR Master Mix | PCR | Thermo Fisher Scientific™ | 4× | |
| TURBO™ DNase | PCR | Thermo Fisher Scientific™ | 2 U/µL | |
| Magnesium Acetate (MgOAc) | RPA | TwistDx™ | 280 mM | TwistAmp® Basic Kit |
| RevertAid Reverse Transcriptase | RPA | Thermo Fisher Scientific™ | 200 U/µL | |
| RNase H | RPA | NEB® | 5 U/µL | |
| RPA Pellets (lyophilized) | RPA | TwistDx™ | N/A | TwistAmp® Basic Kit |
| RPA Primers (forward, reverse) | RPA | Integrated DNA Technologies™ | 5 µM of each primer | *Table S4* for sequences |
| SuperScript IV Reverse Transcriptase | RPA | Thermo Fisher Scientific™ | 200 U/µL | Invitrogen |
| Synthetic DNA Target | RPA | Integrated DNA Technologies™ | $10^{10}$ copies/µL | *Table S4* for sequences |
| Synthetic RNA Target | RPA | N/A | $10^{10}$ copies/µL | generated via *in vitro* transcription of synthetic DNA target |
| Nuclease-free Water | RPA; SHERLOCK | Thermo Fisher Scientific™ | N/A | Invitrogen |
| Reaction Buffer (Optimized) | RPA; SHERLOCK | N/A | 5× | 0.1 M HEPES pH 8.0, 300 mM KCl, 25% PEG-8000 |
| Reaction Buffer (Original) | RPA; SHERLOCK | N/A | 5× | 0.1 M HEPES pH 6.8, 300 mM NaCl, 25% PEG-8000, 25 uM DTT |
| Cas13a crRNA | SHERLOCK | Integrated DNA Technologies™ | 2 µM | *Table S4* for sequences |
| Cleavage Buffer (CB) | SHERLOCK | N/A | 10× | 400 mM Tris pH 7.5, 10 mM DTT |
| FAM Cleavage Reporter (Biotinylated) | SHERLOCK | Integrated DNA Technologies™ | 16 µM | *Table S4* for sequences |
| polyU FAM Cleavage Reporter (Quencher) | SHERLOCK | Integrated DNA Technologies™ | 2 µM | *Table S4* for sequences |

| | | | | |
|---|---|---|---|---|
| *Lwa*Cas13a protein | SHERLOCK | GenScript® | 0.5 mg/mL | Custom protein purification as described previously (23) |
| Magnesium Chloride (MgCl2) | SHERLOCK | Thermo Fisher Scientific™ | 1 M | Invitrogen |
| HybriDetect Assay Buffer | SHERLOCK | Milenia® Biotec | | |
| HybriDetect 1 Lateral Flow Strips | SHERLOCK | Milenia® Biotec | N/A | |
| RNaseAlert® Substrate v2 | SHERLOCK | Thermo Fisher Scientific™ | 2 μM | |
| rNTPs | SHERLOCK | NEB® | 25 mM of each nucleotide | |
| Storage Buffer (SB) | SHERLOCK | N/A | 1× | 50 mM Tris pH 7.5, 600 mM NaCl, 5% glycerol, 2 mM DTT |
| T7 RNA Polymerase | SHERLOCK | Lucigen® | 50 U/μL | NextGen® |

**Table S4: Oligonucleotides used in this study.**

| Reagent | Sequence | SARS-CoV-2 Gene Location |
|---|---|---|
| Cas13a crRNA | CUCUUCUUCAGGUUGAAGAGCAGCAGAA | Orf1ab |
| PCR Primer (forward) | GACCCCAAAATCAGCGAAAT | N1 |
| PCR Primer (reverse) | TCTGGTTACTGCCAGTTGAATCTG | N1 |
| PCR Probe | /FAM/ACCCCGCATTACGTTTGGTGGACC/ BHQ1 | |
| RPA Primer (forward) | CCAAGGTAAACCTTTGGAATTTGGTGCCAC | Orf1ab |
| RPA Primer (reverse) | ACTATCATCATCTAACCAATCTTCTTCTTG | Orf1ab |
| Synthetic DNA Target | GTGAGTTTAAATTGGCTTCACATATGTATT GTTCTTTCTACCCTCCAGATGAGGATGAAG AAGAAGGTGATTGTGAAGAAGAAGAGTTTG AGCCATCAACTCAATATGAGTATGGTACTG AAGATGATTACCAAGGTAAACCTTTGGAAT TTGGTGCCACTTCTGCTGCTCTTCAACCTG AAGAAGAGCAAGAAGAAGATTGGTTAGATG ATGATAGTCAACAAACTGTTGGTCAACAAG ACGGCAGTGAGGACAATCAGACAACTACTA TTCAAACAATTGTTGAGGTTCAACCTCAAT TAGAGATGGAACTTACACCAGTTGTTCAGA CTATTGAAGTGAATAGTTTTAGTGGTTATT TAAAACTTACTGACAATGTATACATTAAAA ATGCAGACATTGTGGAAGAAGCTAAAAAGG TAAAACCAACAGTGGTTGTTAATGCAGCCA ATGTTTACCTTAAACATGGAGGAGG | Orf1ab |
| T7 Promoter ssDNA Primer | GAAATTAATACGACTCACTATAGGG | dsDNA appended upstream of target |
| FAM Cleavage Reporter (Biotinylated) | /56-FAM/rUrUrUrUrUrUrUrUrUrUrUrU rU/3Bio/ | |
| polyU (*i.e.*, 6U or 7U) FAM Cleavage Reporter (Quencher) | /56-FAM/rUrUrUrUrUrU(rU)/3IABkFQ/ | |
