## Supplementary Protocol for "Integrated sample inactivation, amplification, and Cas13-based detection of SARS-CoV-2"

**Supplementary Protocol:** Example protocol for in-tube fluorescent readout of *N* clinical samples with 20 uL reaction volumes.

A. Nuclease inactivation and viral particle lysis

1. Mix 2.28 * *N* µL TCEP-HCl (0.5 M) with 0.023 * *N* µL EDTA (0.5 M) at room temperature.
2. Add 8.8 µL of each sample [*e.g.*, nasopharyngeal swab in universal viral transport medium (UTM) or saliva] to a strip tube. Add 2 µL TCEP/EDTA mixture and 0.2 µL RNase inhibitor (40 U/uL) to each sample.
3. Incubate samples for 5 minutes at 40 ºC, followed by 5 minutes at 70 ºC (if UTM) or 5 minutes at 95 ºC (if saliva).

B. Viral RNA amplification and detection

For *N* samples with 20 µL reaction volumes and a 6.5% pipetting loss, the reaction factor *M* is *N*/5, rounded up to the nearest integer.

| **Reagent** | **Amount (µL)** |
|---|---|
| RPA Pellets | 1 * *M* |
| Reaction Buffer: 0.1 M HEPES pH 8.0, 300 mM KCl, 25% PEG-8000 (Optimized) (5X) | 21.50 * *M* |
| Cas13a resuspended in SB (50 mM Tris pH 7.5, 600 mM NaCl, 5% glycerol, 2 mM DTT) (2.26 µM) | 2.14 * *M* |
| 6U FAM Reporter (2 µM) | 3.19 * *M* |
| RNase Inhibitor (40 U/µL) | 2.69 * *M* |
| rNTPs (25 mM each nucleotide) | 8.60 * *M* |
| SuperScript IV Reverse Transcriptase (200 U/µL) | 1.08 * *M* |
| RNase H (5 U/µL) | 2.15 * *M* |
| T7 RNA Polymerase (50 U/µL) | 2.15 * *M* |
| Nuclease-free H2O | 50.60 * *M* |
| RPA Primers (5 µM each primer) | 2.45 * *M* |
| Cas13a crRNA (2 µM) | 0.46 * *M* |
| Magnesium Acetate (280 µM) | 5.1 * *M* |

1. Make a master mix of all components listed above, except the RPA pellets and the magnesium acetate. Keep the master mix on ice.
2. Resuspend the RPA pellets with 65 uL of the master mix each, returning the resuspended material to the master mix and mixing well.
3. Add magnesium acetate to the reaction mixture.
4. Aliquot 19 uL master mix into wells of a strip tube that is pre-chilled on ice. Add 1 uL of sample (*i.e.*, HUDSON product) or negative control (*e.g.*, nuclease-free water) to each aliquot, mixing thoroughly.
5. Incubate samples at 37 ºC for 30 to 90 minutes.
6. Download the companion smartphone application, which is available at https://github.com/sameeds/sherlock-reader-app. Follow instructions in the application to upload a picture of the results. The application will return binary outcomes (*i.e.*, whether each sample is positive or negative relative to the negative control).
